# Comparative metatranscriptomics of periodontitis supports a common polymicrobial shift in metabolic function and identifies novel putative disease-associated ncRNAs

**DOI:** 10.1101/847889

**Authors:** Nikhil Ram-Mohan, Michelle M. Meyer

**Affiliations:** Department of Biology, Boston College, 140 Commonwealth Ave. Chestnut Hill, MA 02467, USA

**Keywords:** periodontitis, oral microbiome, metatranscriptomics, non-coding RNA, riboswitch, antisense

## Abstract

Periodontitis is an inflammatory disease that deteriorates bone supporting teeth afflicting ∼743 million people worldwide. Bacterial communities associated with disease have been classified into red, orange, purple, blue, green, and yellow complexes based on their roles in the periodontal pocket. Previous metagenomic and metatranscriptomics analyses suggest a common shift in metabolic signatures in disease vs. healthy communities with up-regulated processes including pyruvate fermentation, histidine degradation, amino acid metabolism, TonB-dependent receptors. In this work, we examine existing metatranscriptome datasets to identify the commonly differentially expressed transcripts and potential underlying RNA regulatory mechanisms behind the metabolic shifts. Raw RNA-seq reads from three studies (including 49 healthy and 48 periodontitis samples) were assembled into transcripts de novo. Analyses revealed 859 differentially expressed (DE) transcripts, 675 more- and 174 less-expressed. Only ∼20% of the DE transcripts originate from the pathogenic red/orange complexes, and ∼50% originate from organisms unaffiliated with a complex. Comparison of expression profiles revealed variations among disease samples; while specific metabolic processes are commonly up-regulated, the underlying organisms are diverse both within and across disease associated communities. Surveying DE transcripts for known ncRNAs from the Rfam database identified a large number of tRNAs and tmRNAs as well as riboswitches (FMN, glycine, lysine, and SAM) in more prevalent transcripts and the cobalamin riboswitch in both more and less prevalent transcripts. In silico discovery identified many putative ncRNAs in DE transcripts. We report 15 such putative ncRNAs having promising covariation in the predicted secondary structure and interesting genomic context. Seven of these are antisense of ribosomal proteins that are novel and may involve maintaining ribosomal protein stoichiometry during the disease associated metabolic shift. Our findings describe the role of organisms previously unaffiliated with disease and identify the commonality in progression of disease across three metatranscriptomic studies. We find that although the communities are diverse between individuals, the switch in metabolic signatures characteristic of disease is typically achieved through the contributions of several community members. Furthermore, we identify many ncRNAs (both known and putative) which may facilitate the metabolic shifts associated with periodontitis.

## Introduction

Afflicting more than three million individuals in the United States per year, periodontitis is an oral disease characterized by inflammation of the periodontium (resulting from poor oral hygiene) that eventually leads to tooth loss. Given the prevalence of periodontitis, the microbial instigators of the disease have been studied for decades, with novel technologies successively contributing toward a better understanding of the disease and its polymicrobial origins. Cultivation based methods originally identified anaerobic, Gram negative, rods deep in the periodontal pocket (dominated by *Bacteroides melaninogenicus* and *Fusobacterium nucleatum*) (Slots, 1977). Circumventing the limitations of culture-based techniques, 16S rRNA sequencing from the subgingival plaques of 50 individuals with advanced periodontitis revealed eight key periodontal pathogens: *Actinobacillus actinomycetemcomitans, Bacteroides forsythus* (now *Tannerella forsythia*), *Campylobacter rectus, Eikenella corrodens, Porphyromonas gingivalis, Prevotella intermedia, Prevotella nigrescens* and *Treponema denticola* (Ashimoto et al., 1996), and further screening expanded the list of periodontal pathogens to include members of *Deferribacteres, Bacteroidetes*, OP11, and TM7 phyla, and several novel species including *Eubacterium saphenum, Porphyromonas endodontalis, Prevotella denticola*, and *Cryptobacterium curtum* (Kumar et al., 2003).

A subsequent high-throughput DNA hybridization study of periodontitis progression in 185 individuals revealed six complexes of disease associated organisms characteristic of distinct stages of disease progression (red, orange, yellow, green, blue and purple) (Socransky et al., 1998). The red complex, comprised of *Porphyromonas gingivalis, Treponema denticola*, and *Tannerella forsythia*, colonizes the biofilm during late stage periodontitis and is the major pathogenic complex (Holt and Ebersole, 2005; Socransky et al., 1998; Ximénez-Fyvie et al., 2000). The orange complex, which is strongly associated with the red complex, is comprised of a larger number of species including – *Campylobacter gracilis, Campylobacter rectus, Campylobacter showae, Eubacterium nodatum, Fusobacterium nucleatum, Parvimonas micra, Prevotella intermedia, Prevotella nigrescens*, and *Streptococcus constellatus*. As colonization by the orange complex progresses, more members of the red complex also colonize, suggesting that during disease, colonization of the orange complex directly precedes the red complex (Socransky et al., 1998; Socransky and Haffajee, 2005). Members of the other four complexes (purple, blue, yellow, and green) partake in the initial colonization of the periodontal pocket causing a cascading effect leading toward the orange and red complexes (Socransky and Haffajee, 2005).

The advent of metagenomic analysis has further underscored that periodontitis is a result of polymicrobial synergy and dysbiosis (Lamont and Hajishengallis, 2015). The mean species diversity of the microbial community appears to change drastically between healthy and periodontitis affected states. However, the precise effect is not well understood. Initial 454-pyrosequencing of 16S rRNA libraries followed by qPCR of 22 chronic periodontitis samples uncovered higher alpha diversity and biomass associated with the disease community. This finding suggests that new dominant taxa emerge, but the original health-associated community may not be replaced (Abusleme et al., 2013). However, subsequent meta-analysis across several studies shows reduced alpha diversity associated with disease (Ai et al., 2017). Based on fluctuations of relative metagenome abundances there appear to be a set of marker species that differentiate healthy, stable, and progressing sites of periodontitis (Ai et al., 2017). Furthermore, comparison of 16 metagenomic samples revealed the existence of a core disease affiliated community (Wang et al., 2013). However, the keystone pathogens of periodontitis that interfere with host immune defenses leading to tissue destruction (*Porphyromonas gingivalis* (Darveau et al., 2012; Hajishengallis et al., 2011, 2012; Orth et al., 2011), *Prevotella nigrescens*, and *Fusobacterium nucleatum* (Szafrański et al., 2015)) are not among the identified marker species that differentiate diseased and healthy samples. The observed changes in community structure go hand in hand with an alteration in the functional profile of the community. Metagenomic surveys of healthy and diseased dental plaques revealed that genes encoding bacterial chemotaxis, motility, and glycan biosynthesis and metabolism are over-represented in disease whereas metabolism of carbohydrates, amino acids, energy, and lipids, membrane transport, and signal transduction are under-represented (Wang et al., 2013).

Three metatranscriptomic surveys have provided further insight directly into metabolic activity during disease progression (Duran-Pinedo et al., 2014; Jorth et al., 2014; Yost et al., 2015). Gene ontology enrichment analyses of metatranscriptomes from progressing and stable periodontitis sites revealed that members of the red complex up-regulate their TonB-dependent receptors, aerotolerance genes, iron transport genes, hemolysins, and CRISPR-associated genes, and enzymes like proteases and peptidases (Yost et al., 2015). However, transcripts (for processes such as proteolysis, potassium transport, and cobalamin biosynthesis) from organisms not previously associated with disease also show differential expression, suggesting involvement of additional organisms (Yost et al., 2015). Functional comparisons of healthy and aggressive periodontitis sites revealed that upregulation of lysine fermentation, histidine degradation, and pyruvate metabolism are common to the diseased individuals (Jorth et al., 2014). The collective finding of all three metatranscriptomic studies is the conservation of the community functionality rather than the specific microbial effecters of disease (Duran-Pinedo et al., 2015; Jorth et al., 2014; Yost et al., 2015). This finding emphasizes how understanding of the etiology of disease has progressed from the red complex instigators, to the keystone pathogen concept, and to finally toward a polymicrobial synergy and dysbiosis model (Hajishengallis and Lamont, 2012; Lamont and Hajishengallis, 2015).

While most of the emphasis of past metatranscriptomic analysis has been on identifying up-regulated protein coding regions and organisms associated with these coding regions, a functional bacterial transcriptome also includes many non-coding RNAs (ncRNAs). ncRNAs are untranslated, often structured, elements that are key posttranscriptional regulators acting on mRNA degradation (Desnoyers et al., 2013), translation initiation (Frohlich and Vogel, 2009; Urban and Vogel, 2007), synthesis of ribosomal proteins (Deiorio-Haggar et al., 2013), and transcription attenuation (Breaker, 2012) in response to environmental cues. These regulatory elements can be divided into *cis-* and *trans-*acting based on their location on the genome with respect to the regulated target. Riboswitches are classical examples of *cis*-acting ncRNAs that are typically found in the 5’-untranslated regions (5’UTR) immediately upstream of the regulated gene. ncRNAs are also found antisense to coding regions and often interfere with transcription (Neufing et al., 2001) or repress translation (Sayed et al., 2012). Only the metatranscriptomic study conducted by Duran-Pinedo et al. screened for differentially expressed ncRNAs in periodontitis. This study identified 20 Rfam families within a reported 12,097 small RNAs (sRNAs) overrepresented in disease (Duran-Pinedo et al., 2015). Activities regulated by these ncRNAs include: amino acid metabolism, carbohydrate metabolism, control of plasmid copy number, response to stress, and ethanolamine catabolism (Duran-Pinedo et al., 2015). However, Duran-Pinedo et al. defined sRNA very broadly as any transcribed, but not protein-coding, genomic region, and only screened for known ncRNAs (i.e those in the Rfam database (Burge et al., 2013; Daub et al., 2008; Griffiths-Jones et al., 2003, 2005; Nawrocki et al., 2015)) to identify biologically relevant regions. Thus, it is likely that there are additional ncRNAs with biological function in the oral metatranscriptome associated with periodontitis progression that have not been well-described.

Despite decades studying periodontitis and its microbial instigators, the overarching mechanism of disease is still unknown. The organisms, underlying genes, and a short list of ncRNAs driving the disease have been assessed, but the underlying common mechanism of disease progression across the multitude of studies is yet unknown. The current study was undertaken as a meta-analysis to elucidate the genes and potential regulatory mechanisms common to the progression of periodontitis in patients across multiple studies and to illuminate the extent of variation in functional composition between individuals. We achieved this by combining existing RNA-seq read data from three previously published studies (Duran-Pinedo et al., 2014; Jorth et al., 2014; Yost et al., 2015) resulting in a total of 97 pooled metatranscriptome datasets (49 healthy and 48 diseased) to provide increased statistical support and detect subtle differences between the states and the studies (Gibbons et al., 2018). In contrast to previous studies, we also employed a de novo transcript assembly method rather than aligning the reads to a ‘super genome’ of the ∼400 available oral genomes in the Human Oral Microbiome Database (HOMD) or limiting our analysis to only protein coding regions. In addition to screening for known non-coding RNAs, we also engaged a de novo discovery of RNA secondary structures in the assembled transcripts. Our analyses support findings from the earlier studies. We employed standardized cross-sample normalization and identified the common differentially expressed genes and known ncRNAs across 48 disease samples suggesting a shift in metabolic signatures during progression of periodontitis and identified many novel putative structured ncRNAs revealing the potential for riboregulation in periodontal disease.

## Materials and Methods

### Data sources

Previously published RNA-Seq data (Duran-Pinedo et al., 2014; Jorth et al., 2014; Yost et al., 2015) was compiled to result in a total of 49 healthy and 48 disease datasets. Individual fastq files may be found under BioProject accession number SRP033605 as Sequence Read Archives (SRAs) and under submission numbers 20130522 and 20141024 in the publication data repository of HOMD.

### Annotating assembled transcripts

Coding regions within all the Trinity assembled transcripts were identified using TransDecoder (Haas et al., 2013), which was also used for downstream processes such as annotation and binning of the intergenic regions. The assembled transcripts were then annotated two ways. First, the microbial source of each transcript was identified at the species level using BLAST (Altschul et al., 1990) against a local database of the HOMD annotated genomes. Second, the transcripts and identified coding regions were searched against a local Uniprot database using blastx and blastp respectively to determine protein functions. The transcripts were also screened for protein domains using hmmscan (from HMMER (Finn et al., 2011)) against the PFam database (Finn et al., 2014) and rRNA by running RNAMMER (Lagesen et al., 2007). All of the hits were compiled into a SQLite boilerplate database to generate detailed annotation reports for each transcript using Trinotate (Haas et al., 2013). Alongside, the transcripts were also annotated using the KEGG Automatic Annotation Server (KAAS) (Moriya et al., 2007) to derive KEGG Orthology (KO) numbers.

Apart from the functional and taxonomic annotations of the transcripts, we also screened differentially expressed transcripts for known structured non-coding RNA (ncRNA). ncRNAs were identified in the transcripts by searching for every covariance model in Rfam12.2 (Burge et al., 2013; Daub et al., 2008; Griffiths-Jones et al., 2003, 2005; Nawrocki et al., 2015) using cmsearch from infernal-1.1.1 (Nawrocki and Eddy, 2013). Hits with an e value less than 10^-4^ were collected and all non-prokaryotic hits (eg. U1 spliceosomal RNA and Histone3) were dismissed and the remaining ncRNA hits were included in the transcript annotations.

### Differential expression and GO term enrichment analyses

All predicted rRNA transcripts were removed and the expression of the remaining transcripts was quantified in each dataset using Salmon (Patro et al., 2015). Salmon uses a two-phase method employing a quasi-mapping approach as opposed to a traditional alignment-based method to generate count data quickly. A combined matrix was then created for the healthy and disease states with the raw counts for transcript expression. The matrix of expression counts was imported into R (R Core team, 2015) and differential expression analysis was carried out using edgeR (Robinson et al., 2009), the Bioconductor package. Briefly, the counts per million mapped reads (cpm) were calculated for each dataset and low expression transcripts (cpm <0.5) in more than half of the datasets were removed. Since our interests lie in screening for the common effectors in the progression of the disease, we pooled all datasets in each state together (Healthy and Disease) and use each sample as a replicate to estimate common dispersals from the trend in expression. Next, cross sample normalizations using the trimmed mean of M-values (TMM) method was carried out. This normalized expression of transcripts in each sample against an arbitrarily chosen reference sample and excluded outliers. exactTest was run on the filtered transcripts between the healthy and disease states to identify the DE transcripts. Only transcripts with a log fold change (log_2_FC) ≥ 1 or ≤ −1 with a p value < 0.05 were considered significant. Gene Ontology (GO) term enrichment analysis of the DE transcripts was carried out using GOseq (Young et al., 2010), the Bioconductor R package. Gene lengths, of the assembled transcripts, were estimated with a perl script that is a part of the Trinity installation. Enriched and depleted GO terms estimated in health and disease were considered significant if the p value ≤ 0.05. The DE genes were then mapped onto KEGG pathways to precisely locate the steps that were highly up- or down-regulated. These were visualized using Pathview (Luo and Brouwer, 2013, 2015), the Bioconductor package. Further, the major contributing species in each sample for the enriched processes or pathways was identified by surveying the individual metatranscriptome fastq files using HUMAnN2 (Abubucker et al., 2012).

### Taxonomic makeup of disease samples

In order to establish the taxonomic makeup of the disease samples being studied, the metatranscriptomic reads were subject to MetaPhlAn2 (Truong et al., 2015), a part of the HUMAnN2 package for analyzing microbiome datasets (Abubucker et al., 2012). To determine the microbial community composition in each sample we mapped all the reads using Bowtie2 (Langmead et al., 2013) to a custom compiled marker sequence database available with HUMAnN2. Relative abundances were calculated to identify the prevalent species in each sample.

### Comparison of expression profiles between samples

Based on the differential expression analysis, the significant transcripts were selected from the entire transcript collection and the expression profile in each of the 49 healthy and 48 disease samples were compared for these transcripts. Pearson correlation coefficients ranging between −1 for a negative relationship to 1 for a positive relationship were calculated for each pairwise observation using the cor function in R.

### In silico identification of known ncRNAs and de novo discovery of putative structured ncRNA

In an effort to identify ncRNAs that might regulate the progression of periodontitis, the DE transcripts were assayed for both known and novel structured ncRNAs. Known ncRNAs deposited in Rfam12.2 (Burge et al., 2013; Daub et al., 2008; Griffiths-Jones et al., 2003, 2005; Nawrocki et al., 2015) were identified using cmsearch from infernal-1.1.1 (Nawrocki and Eddy, 2013). de novo discovery was performed by GraphClust (Heyne et al., 2012). The intergenic regions were pooled and the nucleotide sequences extracted. The compiled sequences were processed two ways – 1) all of the compiled sequences together; 2) blast hits for the sequences against Refseq77 (O’Leary et al., 2016) were collected and the each set of hits was run separately. GraphClust, employing infernal-1.0.2 (Nawrocki et al., 2009); RNAshapes (Steffen et al., 2006); LOCARNA (Smith et al., 2010; Will et al., 2007, 2012); ViennaRNA (Lorenz et al., 2011); and RNAz (Gruber et al., 2010), was programmed to search for structures in sliding windows of 150 nucleotides with a shift of 75 nucleotides.

The predicted putative structures and their covariance models derived by GraphClust were searched against the entirety of the Refseq77 genomic database using an in house perl pipeline to obtain more, phylogenetically distant hits if possible. In short, the generated covariance model was converted to version 1.1.1 of infernal, calibrated and then searched against the sequence database. High confidence hits were extracted and realigned with the original covariance model. The resultant Stockholm file was then filtered to remove sequences that do not have at least 60% of the predicted structure before manual curation on RALEE (Griffiths-Jones, 2005).

## Results

### Only a fraction of the differentially expressed (DE) transcripts originate from red/orange complexes

RNA-Seq data published by earlier studies (Duran-Pinedo et al., 2014; Jorth et al., 2014; Yost et al., 2015) that surveyed the community composition as well as the functional profile variation between healthy and periodontitis affected sites were collected. Despite many possible differences between the studies – patient profile; disease severity; methodology used; we initially combined the datasets in an effort to identify the underlying commonality across all studies. We accumulated read data from a total of 49 healthy and 48 diseased samples. The collected data were split into the two categories - Healthy with a total of 26,034,228 reads, and Disease with 34,697,369 reads, and concatenated into a combined dataset for de novo assembly using Trinity (Grabherr et al., 2011; Haas et al., 2013). This generated a total of 627,752 transcripts with an average GC content of 49.76%, a median contig length of 298 nucleotides, and an N50 of 420 nucleotides.

Differential expression analyses of the transcripts between healthy and diseased states using edgeR with counts first normalized to each dataset and then cross-sample normalized resulted in a total of 859 DE transcripts with a p value < 0.05 and a log_2_FC ≥ 1 or ≤ −1 (Table S1). Of these, 675 showed increased expression in diseased samples, and 184 displayed decreased expression. Although such differences are the result of both up- or down-regulated gene expression within an organism and frequency changes in members of the microbial community, we will refer to increased prevalence transcripts as up-regulated, and decreased prevalence transcripts as down-regulated throughout this manuscript. The transcripts are represented by 157 species and strains spanning 52 genera. Only ∼20% of the up- or down-regulated transcripts originated from the members of the red or orange complexes and ∼50% originate from microbes unaffiliated with any specific complexes (Fig. 1a). Interestingly, many unaffiliated microbial species are closely related to those previously classified into the microbial complexes and show large counts of DE transcripts. For example, members of the genus *Streptococcus* account for 212 of the 859 DE transcripts and span many of the disease associated complexes. *S. constellatus* is grouped in the orange complex; and *S. gordonii, S. intermedius, S. mitis, S. sanguinis*, and *S. oralis* are classified in the yellow complex. However, *S. anginosus, S. australis, S. cristatus, S. infantarius, S. infantis, S. mutans, S. oligofermentans, S. vestibularis, S, pneumoniae, S. peroris, S. parasanguinis*, and several *Streptococcus oral* taxa are unaffiliated with specific complexes yet still contribute significantly to the DE transcripts.

**Figure 1.**
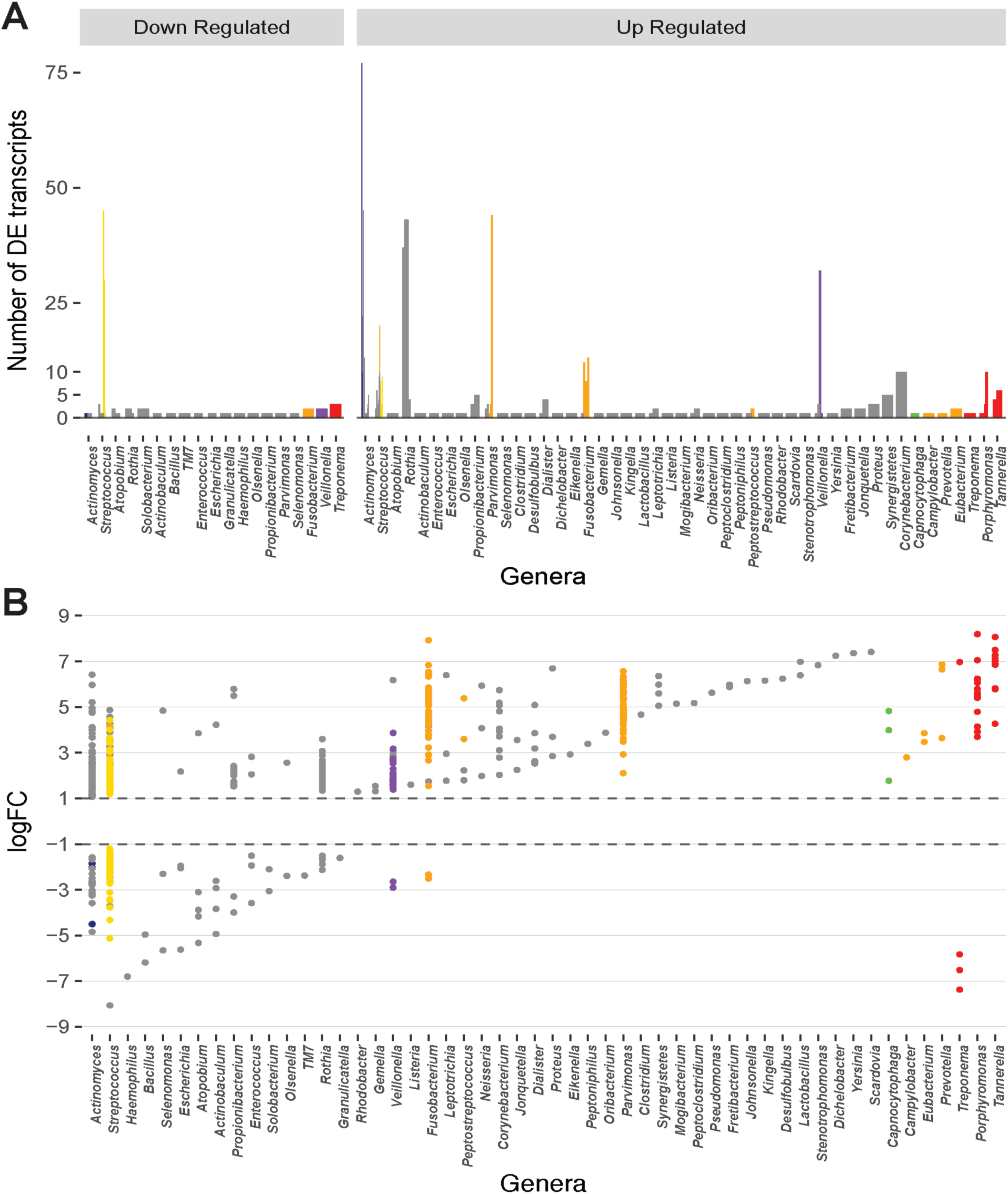
Distribution of the number and magnitude of differentially expressed (DE) transcripts in the genera represented in the oral microbiome. Distribution of the origin of DE transcripts. **A.** The number of DE transcripts originating from different genera. Juxtaposed lines represent different species within a genus that have DE transcripts. Bars are colored based on the classification of the organism into periodontitis associated complexes (Socransky et al., 1998). *Actinomyces* and *Streptococcus* are the genera with the largest number of different species that undergo differential expression during disease. **B.** Magnitude of up- or down-regulation (Log_2_ fold-change) of the transcripts during disease. Each point represents a DE transcript and points are colored by the complex in which the organism is classified (Socransky et al., 1998). A large fraction of the differentially transcripts originate from organisms that are unaffiliated to a specific disease associated complex. Transcripts from members of the red and orange complexes undergo higher magnitude differential expression.

The up- and down-regulated transcripts originate from 121 and 63 bacterial species respectively. The ten species with the largest number of up-regulated transcripts are shown on Table 1, and the majority are not affiliated with a disease associated complex. Members of the orange complex account for ∼18% of the up-regulated transcripts whereas red complex members account for only ∼4%. Of the down-regulated transcripts, ∼24% and 16% are derived from *Streptococcus sanguinis* SK36 and *Streptococcus gordonii Challis* CH1 respectively, both of which are members of the yellow complex. Three transcripts from *Treponema* (red complex), and two from *Fusobacterium* (orange complex) are the only down-regulated transcripts from either complex, consistent with the large presence of these pathogenic bacteria, and the resulting increase in their transcriptome, in diseased samples.

**Table 1.** Species with the largest number of DE transcripts. List of top 10 species with the most number of up-regulated transcripts. Includes information of the microbial complexes these organisms belong to.

| Organism | Complex | % of total up-regulated transcripts |
| --- | --- | --- |
| *Actinomyces viscosus* C505 | Blue | ~11% |
| *Actinomyces* oral taxon 175 | Unaffiliated | ~7% |
| *Parvimonas micra* ATCC 33270 | Orange | ~7% |
| *Rothia dentocariosa* M567 | Unaffiliated | ~6% |
| *Rothia dentocariosa* ATCC 17931 | Unaffiliated | ~5% |
| *Veillonella parvula* DSM 2008 | Purple | ~5% |
| *Actinomyces* oral taxon 171 | Unaffiliated | ~3% |
| *Streptococcus constellatus pharyngis* SK1060 | Orange | ~3% |
| *Fusobacterium nucleatum vincentii* ATCC 49256 | Orange | ~2% |
| *Actinomyces oris* K20 | Unaffiliated | ~2% |

### Differentially expressed transcripts from the orange and red complexes show the greatest magnitude changes

Although the red and orange complexes account for only a small percentage of the differentially expressed genes, the magnitude of change in expression in these transcripts is drastic (Fig. 1b). The up-regulated red complex transcripts range from a log_2_FC of 3.6 to 8.2 with an average expression change of ∼6.1-fold, and the down-regulated transcripts range from −5.8 to - 7.4-fold with an average drop in expression of ∼6.6-fold. Similarly, the change in expression of the up-regulated transcripts from the orange complex ranges from ∼1.5 to ∼7.9-fold with an average change in expression of ∼4.6-fold. Only two transcripts from the orange complex are down regulated with changes in expression of −2.5 and −2.3-fold. Members of the other complexes do not undergo such a drastic change in expression. Change in the expression of transcripts from the blue complex ranges from about 1.2 to 4.8-fold (average: ∼1.9) and −1.9 to −4.5-fold (average: - 2.7); green complex ranges from 1.8 to 4.8-fold (average: 3.5), with no down-regulated transcripts; purple complex ranges from 1.4 to 3.9-fold (average: 2.0), and two down-regulated transcripts with log_2_FCs of −2.9 and −2.6 respectively; yellow complex ranges from ∼1.2 to 4.4 (average: 2.1), and ∼ −1.2 to −5.1 (average: −2.03). Finally, the unaffiliated group, with many diverse genera, displays changes in expression between ∼1.1 to 7.4-fold with an average change in expression of 2.6-fold, and ∼ −1.5 to ∼ −8.1 with an average down-regulation of ∼ −3-fold.

### Expression profiles of disease samples are dissimilar

To assess whether the expression profiles of DE transcripts of disease samples are similar to one another, correlation between profiles of individual samples was estimated. The Pearson correlation coefficient (PCC) was calculated for each pairwise comparison of samples. The PCC is a measure of the linear correlation between two variables where the resulting value between −1 and 1 suggests either a negative linear correlation or a positive one respectively. From a heatmap of the correlation values (Fig. 2), it is immediately apparent that there is no strong correlation between all the healthy samples. This is expected as the oral microbiome in each individual is likely to be highly complex and variant. There is also no strong correlation in the expression profiles of the disease samples. In fact, samples tend to cluster based on the study in which they were generated in rather than the disease state. However, intra study clustering is also not absolute, there are instances where samples from different studies cluster together. This finding is consistent with the diverse criteria for sample inclusion, and methodology across the original studies.

**Figure 2.**
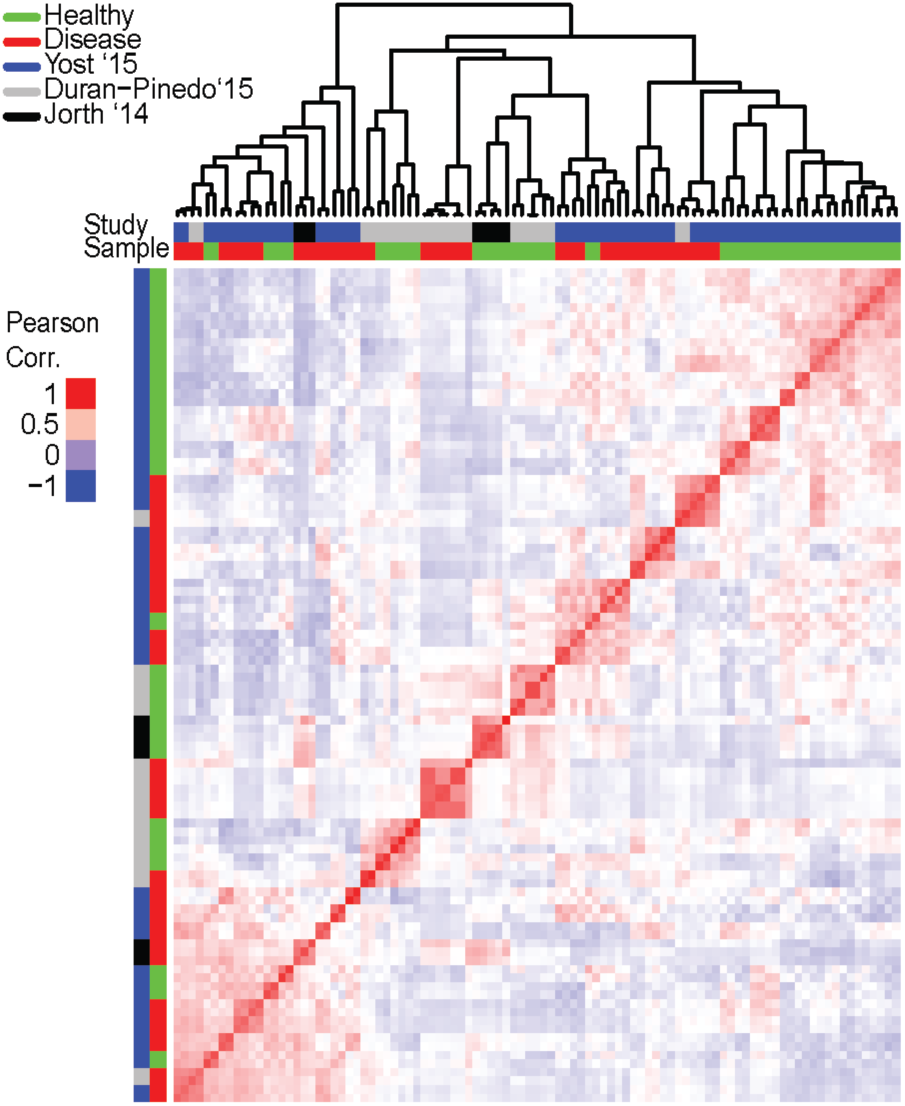
Correlation in expression of differentially expressed (DE) transcripts among individual samples. Expression of the DE transcripts (from the pooled dataset) was calculated for in each individual sample and the pair-wise correlation between samples based on these transcripts computed and a heatmap generated to represent the correlation. Red is high correlation (PCC = 1) while blue is a negative correlation (PCC = −1). Clustering of samples based on the similarity in expression is represented on top (since this is a pairwise comparison, only the column dendrogram is represented). Samples are further annotated by state – disease or healthy, and by origination study. Clustering of disease samples across different origination studies based on expression is not observed. Possible clustering of samples based on the origination study suggests inherent differences between studies and may be attributed to many variables such as inclusion criteria, disease progression, or sampling and sequencing methodologies.

The lack of correlation in the expression profiles can be attributed to the variation in the community composition of the samples. The genus and species level composition of each disease sample (using the entire metatranscriptomes) show distinct communities in each sample (Fig. 3a). The differences are apparent not only in variations in the relative abundances of members between communities, but also in the presence/absence of members of each genus and species. Assessing the top 50 most abundant genera in the entire dataset (Fig. 3a), only 17 genera are commonly found in >50% of the samples (Fig. 3b). Members of the *Streptococcus* genus were most frequently detected (∼96% of the disease samples). Members of genera *Rothia* and *Veillonella* were the next commonly detected bacteria in the disease samples (∼92%). Despite their strong association with pathogenicity, members of the *Porphyromonas, Tannerella, Fusobacterium*, and *Treponema* genera were only detected in ∼65%, ∼56%, ∼52%, and ∼52% of the disease samples respectively. Similarly, at the species level, of the 100 most abundant species in the entire dataset, only 14 species were found in >50% of the disease samples. Unlike the genus level, no single *Streptococcus* species is detected in a majority of the samples. However, *Rothia dentocariosa, Actinomyces viscosus, Porphyromonas gingivalis, Tannerella forsythia, Veilonella parvula*, and *Fusobacterium nucleatum* are well represented. The analyses also find highly active viruses in the periodontal pockets including *Tobamovirus, Endornavirus, Gammaretrovirus, Potyvirus*, and *Mastadenovirus.* Thus, the discrepancy in correlation of expression of DE transcripts between disease samples is likely a result of the drastic variation in the functionally active taxa among samples, and is not likely purely due to differences between originating studies.

**Figure 3.**
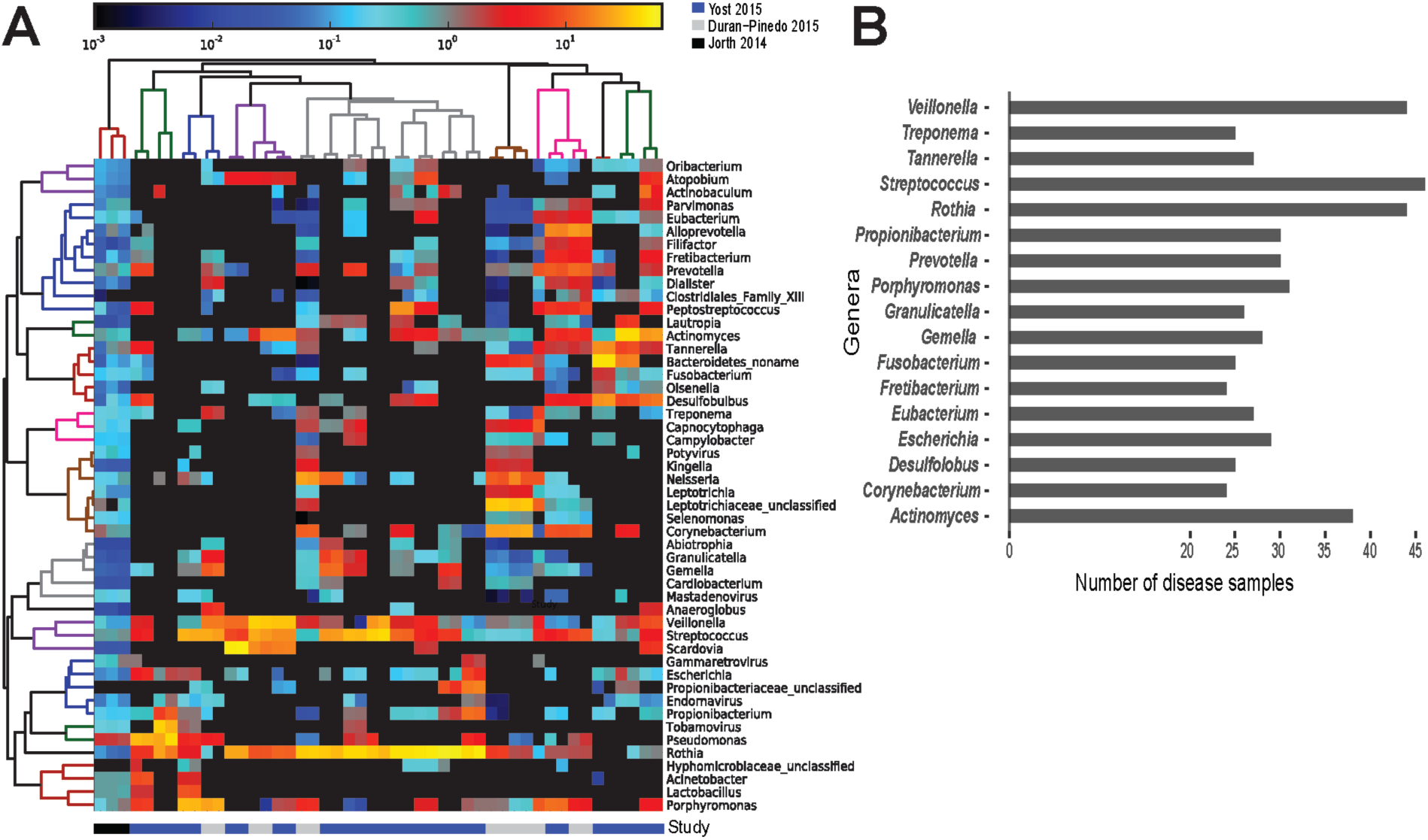
Prevalence of functionally active taxa across disease samples. Screening for the functionally active microorganisms in the disease samples. **A.** Heatmap of the 50 most functionally active genera from the overall dataset identified from the metatranscriptomes using HUMAnN2 (Abubucker et al., 2012). Gradient of color represents the abundance of the genus in a disease sample. Dendrograms represent hierarchical clustering of samples (column) and genera (rows) representing the similarity in the taxonomic composition between the samples and relative abundance of various genera respectively **B.** Frequency distribution of the genera identified in at least 24 of the 48 disease samples. *Streptococcus, Rothia*, and *Veillonella* were found active in 40 out of the 48 disease samples. *Porphyromonas, Tannerella*, and *Fusobacterium* were found active in 30 samples or fewer.

### Specific metabolic processes are enriched in disease

To identify the biological processes that are enriched in the DE transcripts, GO term enrichment analyses using GOseq was conducted. This analysis revealed that biological processes that involve only one organism are significantly less expressed in disease including lipid metabolic processes, hydrolase activity, and peptidase activity in individual organisms. The analyses also revealed an enrichment of genes categorized under the umbrella biological process of localization (e.g. transporter proteins, protein localization to cell surface). These are likely involved in establishment of pathogen localization in the pocket, as periodontitis progresses by the successive localization of various microorganisms in the pocket, until the members of the red complex arrive to drive pathogenesis. Other enriched processes include transport (cation, organic substance, nitrogen compounds, proteins, amino acids), biosynthesis (nitrogen compounds, aromatic compounds, and RNA), metabolism (catabolism of organic substances, glycolytic processes, pyruvate and amino acid metabolism), transcription, and translation.

### Metatranscriptomics supports the polymicrobial nature of periodontitis

The polymicrobial nature of a periodontal infection is readily apparent in the diversity of microorganisms observed to be the sources of the DE transcripts. Mapping the DE transcripts on to KEGG pathways using Pathview revealed communities working in unison to provide a functional shift in disease. For example, mapping the increased prevalence transcripts on to the KEGG pathway for pyruvate metabolism (ko00620) showed both strongly and weakly increased expression genes in the pathway. Surprisingly, annotating these transcripts by their microbial sources revealed DE transcripts originating from members of several complexes (Fig. 4). Different steps along the pathway were increased or decreased prevalence in distinct organisms within the community.

**Figure 4.**
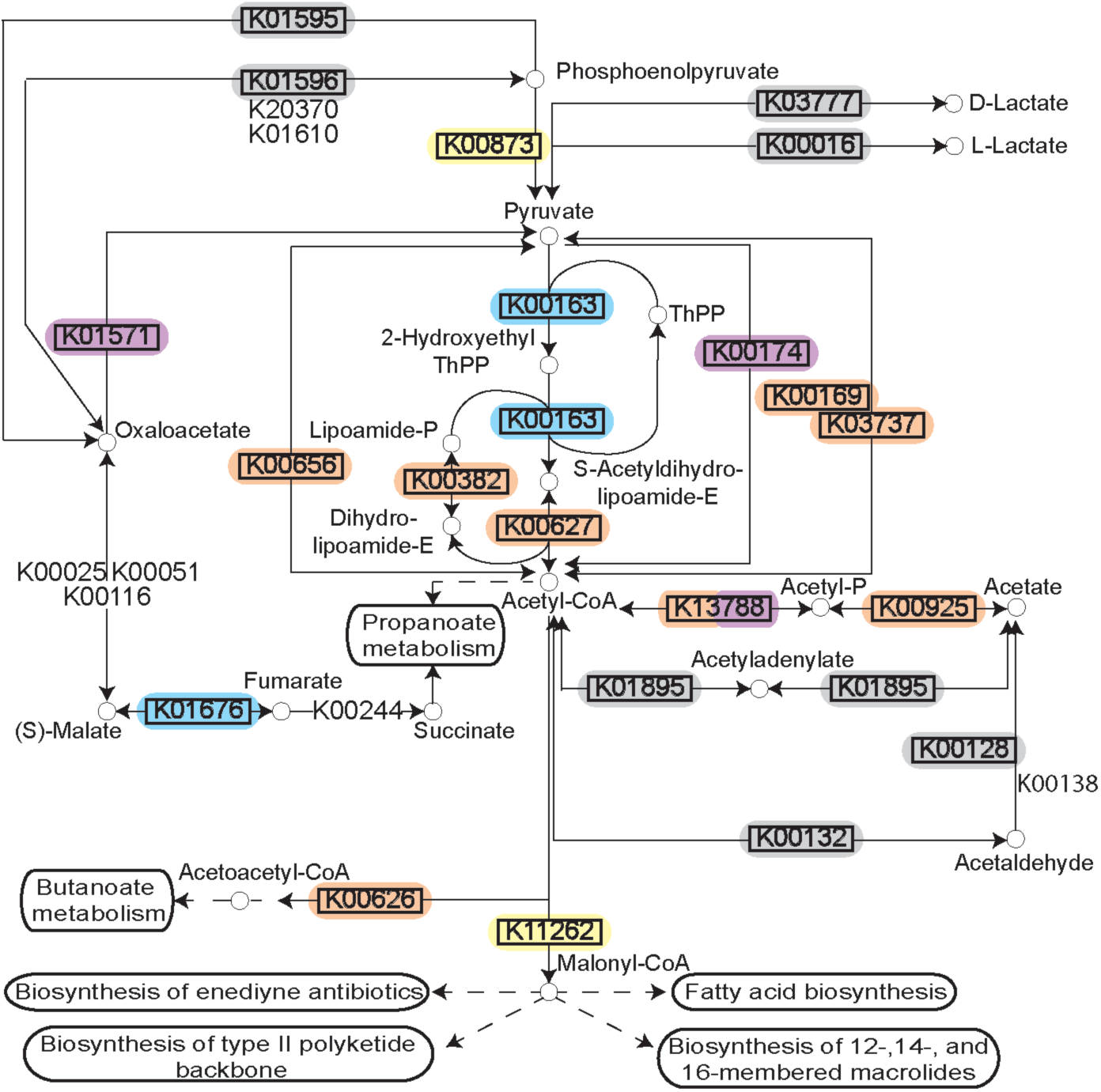
Pyruvate metabolism during disease. Rendering of the KEGG pathway of pyruvate metabolism annotated with the DE transcripts and the microbial complexes from which they originate. Differential expression is represented by the black rectangle around the KEGG Orthology term. Background color for the KEGG orthology term represents the microbial complex of the organism that showed differential expression. The pathway as a whole is up-regulated, but the individual genes are up-regulated in diverse species supporting polymicrobial synergy during periodontitis.

To further assess this observation, the microbial contributors to the pyruvate pathway in each individual disease sample were identified using HUMAnN2. The variation observed in the species contributing to pyruvate metabolism in different samples is striking (Fig. 5a&b). Up-regulation of genes involved in the fermentation of pyruvate to acetate and lactate were detected in 38/48 disease samples (Fig. 5a). Other than four samples where *Streptococcus tigurinus* and two samples where *Rothia aeria* are the sole contributors in the pathway, the remaining 44 samples show a combination of various species contributing to the pathway, a community effort. Similarly, genes involved in the fermentation of pyruvate to isobutanol were up-regulated in 46/48 disease samples (Fig. 5b). Again, other than the 8 samples that display a single organism as the sole contributor to this pathway - *Streptococcus tigurinus* (2 samples), *Streptococcus mitis* (2 samples), *Rothia dentocariosa* (4 samples), the remaining show a combination of organisms contributing to the pathway. This phenomenon is not unique to the pyruvate metabolism pathways. Similar patterns emerge when mapping the DE transcripts to the glycolysis/gluconeogenesis pathway, TCA cycle, lysine biosynthesis pathway, and butanoate metabolism to list a few. In contrast, upregulation of the one carbon pool by folate pathway is driven solely by the members of the orange complex, the zinc/manganese/iron transporters are up-regulated only in the members of the orange complex, and the fermentation of lysine to butanoate is achieved only by *Fusobacterium nucleatum* and *Fusobacterium periodonticum* in 17/48 disease samples. Nonetheless, most metabolic pathways that are enriched under disease conditions seem to have multiple contributing species fulfilling the metabolic niche within a single diseased sample, and different steps along the pathway show increased prevalence due to various contributory species.

**Figure 5.**
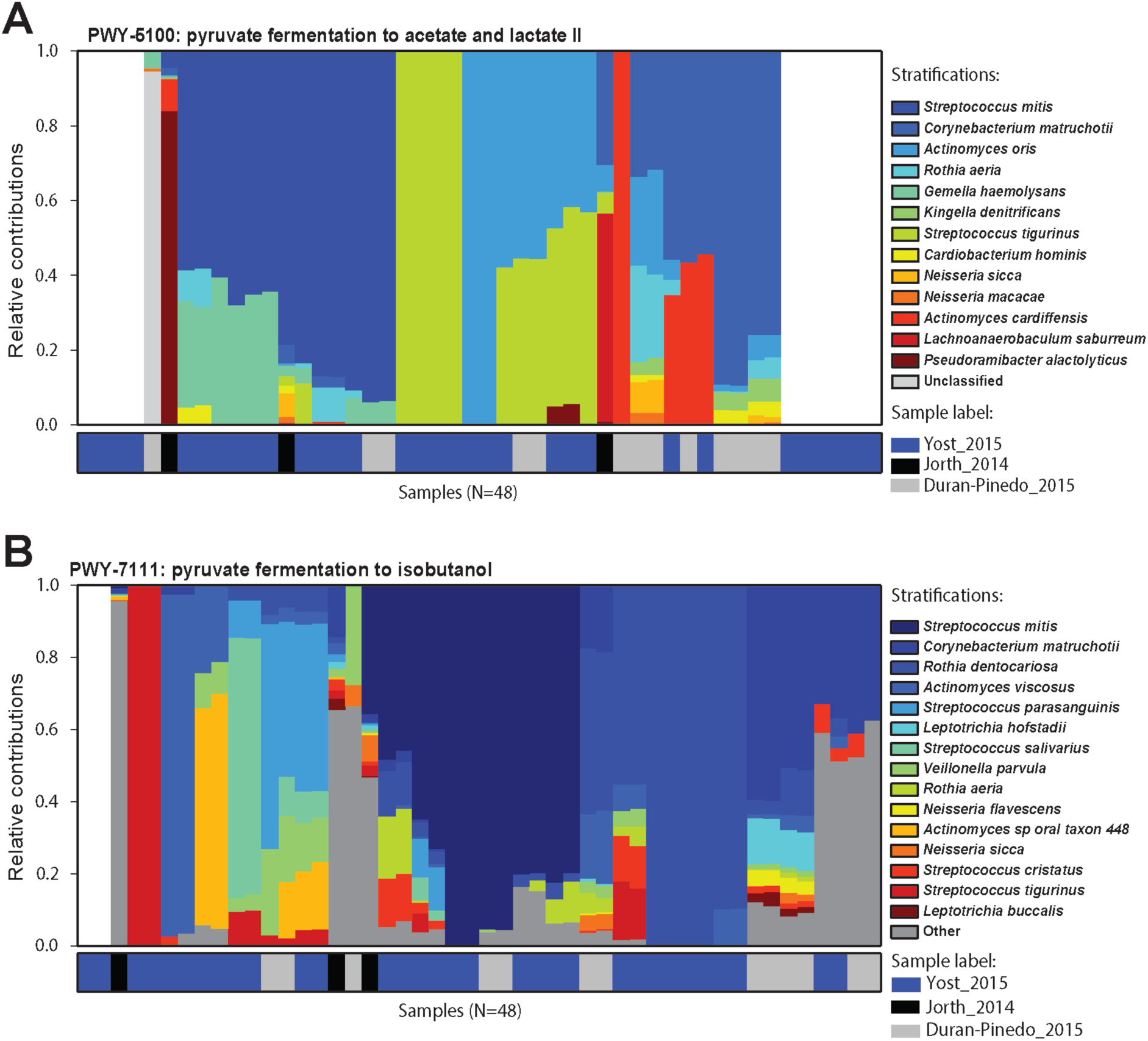
Contributors to pyruvate metabolism during disease. Relative contributions of different bacteria in the two pyruvate metabolism pathways in different samples. **A.** Relative contributions of various organisms in the fermentation of pyruvate to acetate and lactate in each disease sample. Top 15 contributing organisms are listed by name and the rest are grouped as others. Columns are annotated by the study they come from. Different organisms perform similar functions in different disease samples maintaining the functional signature despite the community composition. **B.** Relative contribution of various organisms in the fermentation of pyruvate to isobutanol. Again, functional signature is maintained despite differences in the organisms carrying out fermentation in the different samples.

### Known structured non-coding RNA are present in the DE transcripts

In an effort to identify possible regulatory mechanisms associated with disease, a screen for known ncRNAs in the DE transcripts was carried out using Infernal 1.1.1. This screen returned a total of 635 hits. All non-prokaryotic hits (eg. U1 spliceosomal RNA, Histone3) were removed leaving 181 known structured ncRNAs in the assembled transcripts (Fig. 6). The two predominant classes of ncRNA identified are tRNAs and tmRNAs, which constitute 41% and 19% of the identified ncRNAs identified. These are critical RNA molecules responsible for essential bacterial function, and display both up- and down-regulation depending on the organism. These differences may reflect a change in community members rather than a change in functionality of the community. In addition, the 6S RNA (constituting 3% of the ncRNAs identified) is up- or down-regulated in various unaffiliated genera, and up-regulated in a member of the orange complex – the genus *Eubacterium.* There were also ncRNAs displaying only increased prevalence. RNase P bacterial classes A and B together represent ∼9% of DE ncRNAs. Class A RNaseP originates from members of the blue, orange, and red complexes and unaffiliated organisms, whereas class B only originates from unaffiliated members of genus *Streptococcus*, consistent with the phylogenetic distribution of the two RNase P classes. The bacterial small signal recognition particle (SRP) RNA (∼3% of DE ncRNAs) is also only up-regulated, predominantly by members of the orange complex, namely – *Fusobacterium, Parvimonas*, and *Peptostreptococcus*.

**Figure 6.**
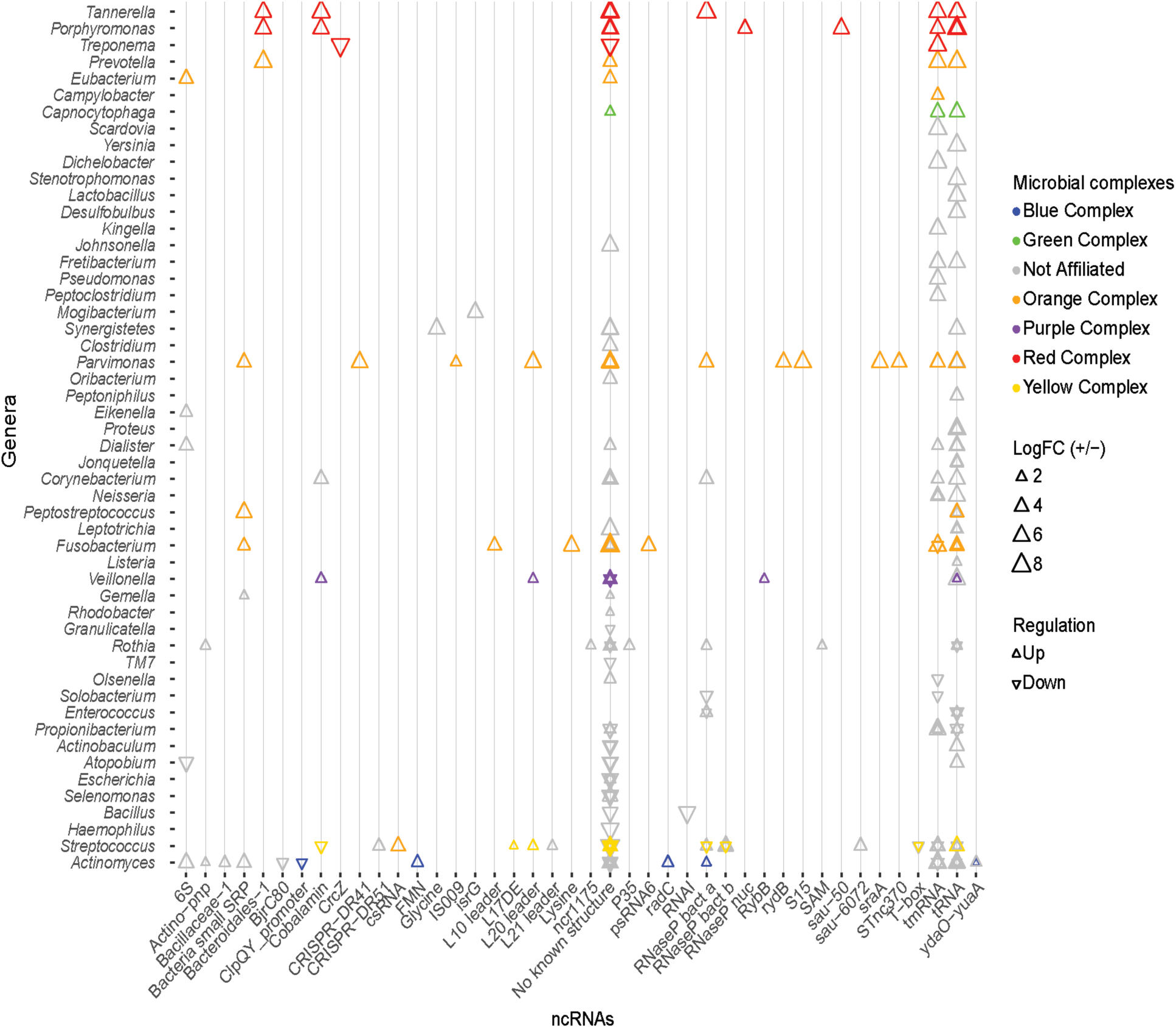
Known ncRNAs identified in differentially expressed transcripts. Known ncRNA identified in DE transcripts, and the genus from which the ncRNA originates. Triangles colors correspond to the organism’s microbial complex, orientation depicts whether the transcript was up- or down-regulated, and the size indicates the magnitude of log_2_ fold change. tRNA and tmRNA are the most abundantly identified ncRNAs. No known ncRNAs were identified in a large number of DE transcripts.

The cobalamin riboswitch is the most predominant regulatory RNA identified (3% of DE ncRNAs). Transcripts containing the cobalamin riboswitch from *Tannerella* and *Porphyromonas* (red complex), *Veillonella* (purple complex), and *Corynebacterium* (unaffiliated) are up-regulated, and transcripts from a species of *Streptococcus* of the yellow complex are down-regulated. These cobalamin riboswitches are associated with different genes in the different organisms. The riboswitch is found upstream of the 4-hydroxybutyryl-CoA dehydratase gene (adenosylcobalamin biosynthesis) in *Tannerella*; upstream of a hypothetical protein in *Porphyromonas*; upstream of methylmalonyl-CoA mutase large subunit gene (adenosylcobalamin dependent enzyme) in *Veillonella*; upstream of a transferase gene in *Corynebacterium*; and upstream of the ATP:cobalamin adenosyltransferase gene (adenosylcobalamin biosynthesis) in *Streptococcus.* Other riboswitches identified in the DE transcripts include the FMN, glycine, lysine, SAM, and the cyclic di-AMP riboswitch (Nelson et al., 2013). Apart from the ncRNAs listed above, many bacterial small RNAs (sRNAs) were also identified from a variety of genera in very low quantities. Various ribosomal protein regulatory elements were also more prevalent in the disease associated transcripts. These include the L10 leader from *Fusobacterium sp.*; the L17 downstream element from a yellow complex member of the genus *Streptococcus*; the L21 leader in an unaffiliated member of the *Streptococcus* genus; the S15 leader in *Parvimonas sp.*; and the L20 leader in *Parvimonas sp., Veillonella sp.*, and a yellow complex member of *Streptococcus.* Although a variety of known ncRNAs including riboswitches and small RNAs were identified in the DE transcripts, the majority of DE transcripts display no secondary structures of known function.

### Novel putative non-coding RNA in the DE transcripts

The large number of DE transcripts containing no previously described secondary structured RNA (Fig. 6) were analyzed using GraphClust to discover novel putative ncRNA structures. This resulted in a total of 224 putative ncRNAs in the up-regulated transcripts and 126 ncRNAs in the down-regulated transcripts. Each of these was manually curated to remove structures with minimal covariation in the predicted base pairing, or lacking a defined genomic context. Of these ncRNAs, 9 putative ncRNAs from the up-regulated and 6 putative ncRNAs from the down-regulated transcripts were scanned against the genomic database Refseq77 using cmsearch (Infernal 1.1.1) to identify additional homologs and determine the phylogenetic distribution of the putative regulatory element. Alignments for each of the 15 putative ncRNAs were analyzed using R-scape (Rivas et al., 2016a, 2016b) to estimate statistical support for the predicted base pairs. Although the most of the putative ncRNAs were identified upstream of the same gene across taxa, like dihydroxyacetone kinase and the nitrogen fixation gene, many were found to be antisense upstream or downstream of the putatively regulated gene. A subset of the identified putative novel ncRNAs and their phylogenetic distribution are described below (Fig. 7).

Of the 224-predicted putative ncRNAs in the up-regulated transcripts, 9 sense and antisense putative ncRNAs were chosen (Fig. 7a) based on their secondary structure and genomic context (supplemental file). Three of these ncRNAs appear to act as 5’-UTR *cis* regulators. The first, ncRNA-161, was identified in a transcript from *Streptococcus anginosus* CCUG 39159 (logFC of ∼2.9). This ncRNA is in the beginning of dihydroxyacetone kinase and is found only in the Firmicutes (Fig. 7a&c), almost exclusively in *Streptococcus* with the exception of *Bacillus sp.* 1NLA3E. Our second candidate ncRNA-116, was found upstream of the *rnfA* gene in *Fusobacterium nucleatum* ATCC 25586. This nitrogen fixation gene involved in electron transport to nitrogenase (Schmehl et al., 1993) was found to have a fold change of ∼4.3 between healthy and disease samples. Furthermore, the putative riboregulator is more highly distributed, appearing in a diverse group of phyla including *Firmicutes, Proteobacteria, Bacteroidetes, Spirochaetes, Thermotogae*, and *Fusobacteria* (Fig. 7c). The third candidate, ncRNA-66, was identified upstream of a hypothetical protein in *Actinomyces oral taxon* 175 F0384 that had a logFC of ∼1.42 and upstream of a hypothetical protein in *Rothia dentocariosa* ATCC 17931 with a logFC of ∼1.41. Survey of the phylogenetic distribution revealed that ncRNA-66 was found distributed upstream of NAD-dependent dehydrolase, UDP-glucose-4-epimerase, and hypothetical proteins across the *Actinobacteria* and *Spirochaetes* (Fig. 7c).

**Figure 7.**
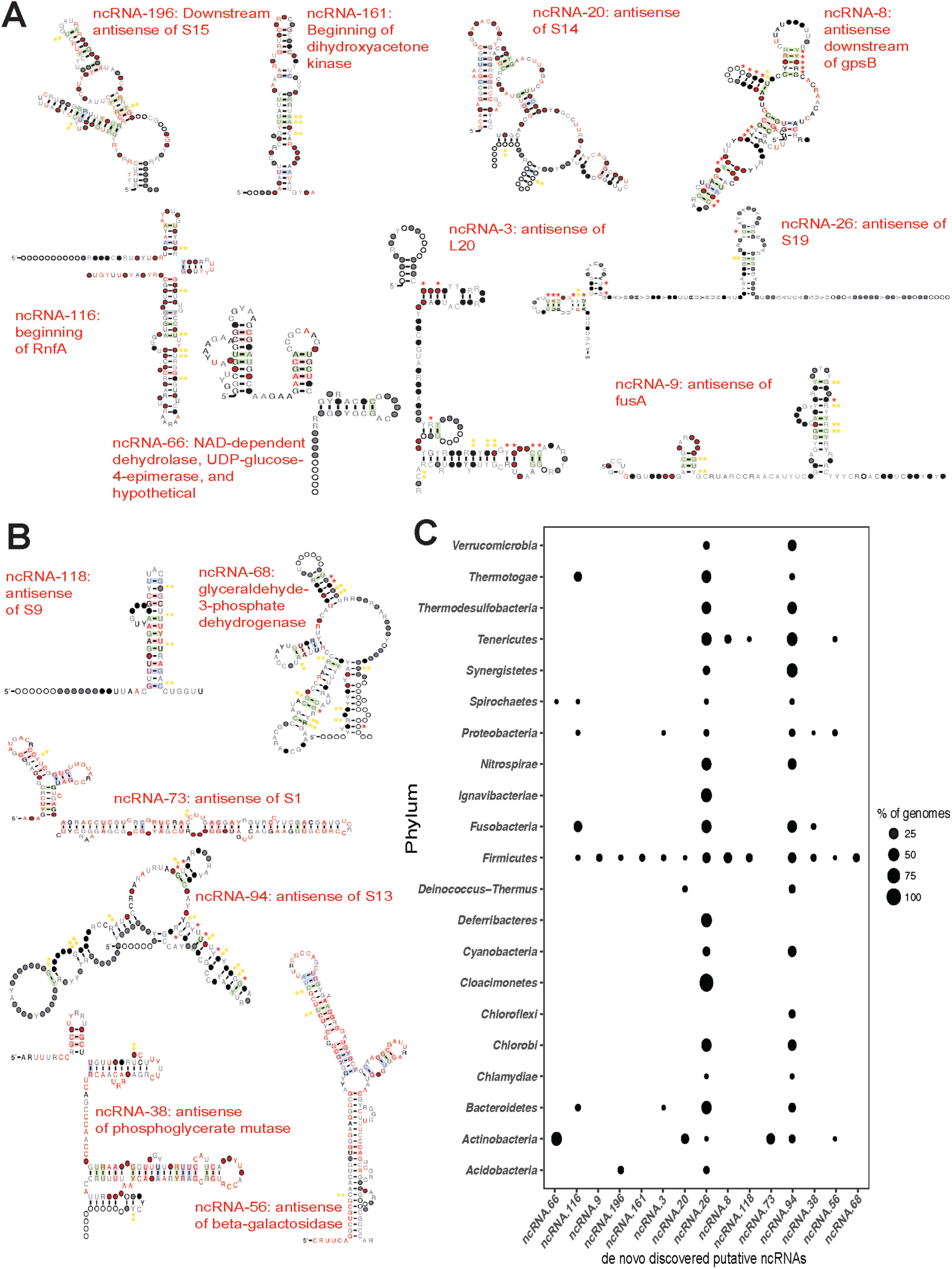
de novo discovered ncRNAs in differentially expressed transcripts. Transcripts that showed no known ncRNAs were subjected to de novo RNA structure discovery using Graphclust (Heyne et al., 2012). Potential ncRNAs were manually curated and a subset of ncRNA were chosen based on predicted secondary structure and evidence for covariation in the base pairing and/or conserved genomic context. Significance of the covariation in predicted base pairs was further tested using R-scape (Rivas et al., 2016a, 2016b). Red * indicate base pairs with E ≤ 0.05 and yellow ** indicate base pairs with E ≤ 1. **A.** Novel putative ncRNAs discovered in the up-regulated transcripts. Consensus sequence and secondary structure of each ncRNA after scanning against Refseq77 and removing hits that did not share at least 60% of the secondary structure. Covarying base pairs are shaded. **B.** Novel putative ncRNAs discovered in down-regulated transcripts produced as described above for the up-regulated transcripts. **C.** Phylogenetic distribution of novel putative ncRNAs in the Bacterial domain. The points represent the percentage of genomes in which the ncRNA was found compared to the genomes in Refseq77 within a phylum. 13 of the 16 novel ncRNAs were found in a fraction of all the Firmicutes genomes. ncRNA-26 (antisense of ribosomal protein S19) and ncRNA-94 (antisense of ribosomal protein S13) are the most widely distributed across the Bacterial domain.

In addition to potential 5’UTR ncRNAs discussed above, our analyses also found four up-regulated putative *cis-*antisense regulators associated with ribosomal protein genes. Our first candidate, ncRNA-196, was identified in three up-regulated transcripts, located downstream and antisense of the ribosomal protein S15 coding region. Two of these transcripts belonged to strains of *Rothia dentocariosa* – M567 and ATCC 17931 with logFC of ∼2.35 and 1.7 respectively, while the third transcript belonged to *Streptococcus oral taxon* 071 73H25AP displayed a logFC of 3.15. This putative ncRNA was found to be distributed through *Firmicutes* and *Actinobacteria* – mostly in the genus *Streptococcus* and in *Bacillus megaterium* WSH-002 within the Firmicutes; and *Rothia dentocariosa* in the *Actinobacteria*. Our second candidate, ncRNA-20 is another example of a putative *cis* antisense regulator of a ribosomal protein, S14. It was found in four up-regulated transcripts, three of which were from the genus *Actinomyces* and one from *Rothia*; *Actinomyces oral taxon* 175 F0384 displayed a logFC of ∼2.21; *Actinomyces oris* K20: ∼1.5; and *Actinomyces naeslundii* MG1: ∼1.5; *Rothia dentocariosa* M567: ∼1.8. Surveying Refseq77 revealed that this ncRNA is distributed across the *Actinobacteria, Deinococcus*-*Thermus*, and *Firmicutes*. A third candidate, ncRNA-3, is found *cis-*antisense of ribosomal protein L20 and represented in two transcripts, both from the genus *Streptococcus – Streptococcus peroris* ATCC 700780 and *Streptococcus cristatus* ATCC 51100 with a fold change of ∼1.69 and ∼1.66 respectively. Phylogenetic distribution of this ncRNA spans the *Firmicutes, Proteobacteria*, and *Bacteroidetes*. Finally, we identified a putative ncRNA antisense of ribosomal protein S19 (ncRNA-26). This ncRNA was found in three highly up-regulated transcripts from *Tannerella forsythia* (logFC ∼7.5), *Fusobacterium nucleatum* (logFC ∼4.0), and *Veillonella parvula* (logFC ∼2.3). It is widely distributed in the Bacterial domain. Of these antisense ncRNAs for ribosomal proteins, we find additional evidence for expression of ncRNA-196 and ncRNA-3 in our recent study of the *Streptococcus pneumoniae* TIGR4 transcriptional profile (Warrier et al., 2018).

We also find up-regulated putative antisense ncRNAs associated with a variety of other processes. Our first candidate, ncRNA-9 is associated with the elongation factor G gene, *fusA*, that is differentially expressed in *Streptococcus oligofermentans* and *Streptococcus anginosus*. It is narrowly distributed and is identified only in ∼6% of the *Firmicutes* genomes in Refseq77. A second example is ncRNA-8, which is antisense and downstream of the cell cycle protein *gpsB*. We find ncRNA-8 in *Streptococcus infantarius* ATCC BAA-102 with the transcript having a logFC of ∼2.5. ncRNA-8 is narrowly distributed to only *Firmicutes* and *Tenericutes*.

We also identified six promising antisense ncRNAs in the down-regulated transcripts (Fig. 7b&c). Many of these also putatively regulate ribosomal proteins. ncRNA-118 was identified antisense of ribosomal protein S9 in *Streptococcus mutans* UA 159 (down-regulated by ∼-1.48 fold). This putative ncRNA is narrowly distributed across Refseq77 and is identified only in *Firmicutes* and *Tenericutes* (Fig. 7c). A second example is ncRNA-73, which is antisense to the beginning of the ribosomal protein S1 coding region in *Actinomyces* oral taxon 180 F0310 (down-regulated ∼-1.6 fold). Surveying the bacterial genomes in Refseq77 revealed that this ncRNA is unique to the *Actinobacteria*. A third example is the putative ncRNA antisense of ribosomal protein S13, ncRNA-94, which was identified in *Granulicatella adiacens* ATCC 49175 (down-regulated ∼-1.6 fold). ncRNA-94 is widely distributed across the Bacterial domain and is absent only in certain phyla such as *Acidobacteria, Claocimonetes*, and *Deferribacteres.*

In addition to ribosomal proteins, we also find down-regulated transcripts antisense to genes involved in sugar metabolism. ncRNA-56 was identified in the *Klebsiella pneumoniae* located antisense to the beginning of the beta-galactosidase gene, and is extremely down-regulated (−6.8 fold). Surveying its distribution, ncRNA-56 is found in the genomes of other *Actinobacteria, Firmicutes, Proteobacteria*, and *Tenericutes.* We also find two antisense ncRNAs putatively regulating different steps of glycolysis. ncRNA-38 was discovered antisense and overlapping the beginning of the down-regulated phosphoglycerate mutase gene. It was found in a fraction of the *Firmicutes, Fusobacteria*, and *Proteobacteria* (Fig. 7c). The other putative regulator of glycolysis is ncRNA-68, which is found antisense and overlapping the translational start of glyceraldehyde-3-phosphate dehydrogenase. However, ncRNA-68 was found only in the *Firmicutes.* By applying de novo discovery pipelines on the DE transcripts, we have identified several promising sense and antisense putative regulators of bacterial ribosomal proteins and other metabolic genes that are associated with periodontitis as reflected in their expression in the healthy vs. disease samples.

## Discussion

Here we present a meta study of three existing sets of metagenomes and metatranscriptomes (Duran-Pinedo et al., 2014; Jorth et al., 2014; Yost et al., 2015) related to oral health and disease to identify commonality in the progression of periodontitis. Although more commonly performed with multiple metagenomes, combined metatranscriptomics analyses enable us to increase statistical support for the findings that are made, as well as understand the commonalities between different studies. From our analyses we find that nearly ∼50% of DE transcripts were from bacteria not previously classified into disease associated complexes (Fig. 1), and only 20% originate from organisms of the red and orange complexes. This mimics previous findings that showed putative virulence factors were up-regulated in larger numbers of bacteria that did not belong to the red or orange complexes (Yost et al., 2015). However, despite the small number of DE transcripts originating from the members of the red and orange complexes, these transcripts show the greatest magnitude of up- or down-regulation.

Healthy samples do not cluster based on the transcript expression, and this is not surprising since mature communities are extremely diverse and often show variations between individual sites within the oral cavity (Marsh, 2006). However, similar expression patterns are also not observed across the disease samples. This lack in correlation among the disease samples is somewhat surprising since previous work suggested that the disease associated microbiota are more similar than health associated communities (Jorth et al., 2014). In this case such differences may also be attributed to differences in sample inclusion criteria, batch effects, and methodological biases, of individual studies. However, these are unlikely to be the only reasons for such lack in correlation. Meta-analyses of RNA-seq data from four studies comprising of 6-13 tissues each from 11 vertebrate species using similar cross sample normalization methods revealed clustering of samples by tissue rather than study or species (Sudmant et al., 2015). This suggests that true commonality between studies can be inferred by these meta-analyses despite any batch effects and methodological biases that might exist. To support this, when comparing individual datasets, no single organism, even at the genus level, was functionally active in all the disease samples suggesting inherent variability between communities. Our analyses also find highly active viruses in the periodontal pocket (Fig. 3A) of many disease samples. However, all but the *Mastadenovirus* are plant related and may reflect the individual’s diet during the sampling. This observed large variety in the functionally active bacterial community composition supports the polymicrobial nature of periodontitis.

Previous metatranscriptomics analyses identified a metabolic shift in disease associated communities towards increased nucleotide biosynthesis, iron acquisition, cobalamin transport, and fermentation of lysine, histidine, and pyruvate (Jorth et al., 2014; Yost et al., 2015). Our own GO term enrichment analyses from the pooled datasets mimics these findings suggesting that these functional shifts are common during the progression of periodontitis. Further exploration of the differentially expressed steps along the pyruvate metabolism pathways using KEGG Orthology showed that different enzymes are differentially expressed in diverse organisms. Even within individual metatranscriptomes, we identified a large degree of diversity in the organisms contributing to pyruvate fermentation (Fig. 4, Fig. 5a&b). Thus, our analysis supports a model where a variety of different bacteria may drive the metabolic process, supporting the polymicrobial dysbiotic nature of the periodontal disease (Darveau, 2010; Hajishengallis and Lamont, 2012; Rosier et al., 2014; Szafrański et al., 2015). Furthermore, our findings also support the notion that periodontitis occurs despite idiosyncratic differences between individuals as long as the community undergoes a switch in its functional and metabolic signals (Dabdoub et al., 2016).

Bacterial metabolic processes are finely controlled. Bacterial non-coding RNAs (ncRNAs) modulate posttranscriptional gene expression genes in response to environmental cues (Breaker, 2012) using a variety of mechanisms including: impacting mRNA decay rate (Desnoyers et al., 2013), regulating translation initiation (Frohlich and Vogel, 2009; Urban and Vogel, 2007), and biosynthesis of ribosomal proteins (Deiorio-Haggar et al., 2013). Given the switch in the functional and metabolic signal during periodontal disease, we searched for putative ncRNAs to drive such changes. Surveying the DE transcripts for known ncRNAs revealed trends similar to those previously observed (Duran-Pinedo et al., 2015) with tRNAs and tmRNAs the most abundant ncRNAs identified in the DE transcripts (Fig. 6). We also identified several other ncRNAs including 6S, bacterial small signal recognition particle, and the cobalamin, FMN, glycine, lysine, and SAM riboswitches, ribosomal protein leaders, and bacterial RNase P class A and B. However, no known ncRNAs were identified in a large majority of DE transcripts.

De novo discovery of ncRNAs in DE transcripts lacking known ncRNAs revealed many novel, putative structured elements. Manual curation of these revealed promising ncRNAs identified sense and antisense to coding regions. Some of the promising ncRNAs and their phylogenetic distributions are presented in Fig. 7, several of which have many predicted base-pairs with statistical support. We find putative sense ncRNAs upstream or near the 5’-end of metabolic genes and antisense ncRNAs corresponding to metabolic genes and many ribosomal protein genes including S1, S9, S13, S14, S14, S19, and L20. Despite a suite of well-characterized RNA *cis*-regulators for ribosomal protein genes (Babina et al., 2015; Deiorio-Haggar et al., 2013; Fu et al., 2013, 2014; Slinger et al., 2014), remarkably there is a dearth of evidence for wide-spread antisense regulation. Only two examples have been described to our knowledge: a σ^B^ induced (stress-induced) transcript has been identified in *B. subtilis* that is anti-sense to *rpsD*, resulting in downregulation of *rpsD* transcript and presumably S4 expression (Mars et al., 2015), and an antisense ncRNA spanning 14 genes of a ribosomal protein operon protects the transcript by hiding RNase E sites providing the *Prochlorococcus* MED4 RNA an enhanced half-life during phage infection (Stazic et al., 2011). The seven putative antisense ncRNA we discovered are likely novel regulators of bacterial ribosomal proteins that might conform to the aforementioned mechanisms of actions or have completely novel mechanisms but are likely worthy of further study in the future as potential targets for antimicrobials treating or preventing periodontitis.

## Author Contributions

MMM conceived the study and edited the manuscript. NR performed all the analyses and wrote the manuscript. Both authors read and approved the final manuscript.

## Acknowledgements

The authors would like to thank Jon Anthony for the setup of many of the tools used. This work was supported by an NIH grant (R21DE025051) to MMM.

## Figure legends

**Supplemental Table 1. List of differentially expressed transcripts estimated form the pooled dataset spanning 48 healthy and 49 disease samples.** A list of 859 DE transcripts with logFC >1 or logFC <-1 and p <0.05.

**Supplemental files.** Alignment files for the discovered novel putative ncRNAs.

